# Direct Entorhinal Control of CA1 Temporal Coding

**DOI:** 10.1101/2023.05.27.542579

**Authors:** Matteo Guardamagna, Oscar Chadney, Federico Stella, Qiangwei Zhang, Clifford Kentros, Francesco P. Battaglia

**Author notes:** Co-Senior Author (Clifford Kentros, Francesco P. Battaglia). Corresponding author(s). E-mail(s); Contributing authors. These authors contributed equally to this work (Matteo Guardamagna, Oscar Chadney, Federico Stella).

## Abstract

During behavior, hippocampal neurons fire in consistent theta sequences, organized by the theta rhythm, which have been linked to predictive coding of future actions. The mechanisms of sequence generation are yet unclear, but in the hippocampal CA1 subfield, are thought to involve both major input streams into CA1 neurons, from CA3 pyramidal neurons and directly from entorhinal cortex. We disentangled the role of these two afferent input with highly specific optogenetic inhibition limited to the direct entorhinal afferents of CA1, thereby leaving the rest of the hippocampal-entorhinal circuit intact. While CA1 spatial firing properties were largely unaffected, theta phase precession was largely abolished. Surprisingly, while theta phase precession is thought to generate theta sequences, theta sequences were actually strengthened when it was suppressed. These results suggest that sequence generation is internal to the hippocampus, while the entorhinal inputs may act as a supervisory signal driving learning and representational updates.

## Main

Temporal coding at multiple scales, from milliseconds to seconds, is used by the brain to convey information and process parallel computations. Within the hippocampus, “theta sequences” [13, 16] of spikes from cells with consecutive place fields, rapidly sweeping through a theta oscillation cycle (*≈*100 ms), are a prominent expression of temporal coding. Theta sequences have been linked to memory of events taking place at different times, prediction of future occurrences, and action planning [25], functions that require the comparison of previously stored traces with the current stream of information about the external world [7]. The hippocampal sub-field CA1 is an interesting candidate for this operation as it sits at the confluence of two inputs: a tri-synaptic pathway through the CA3 sub-field (from Entorhinal Cortex Layer II, EC LII) [36] and a direct extra-hippocampal pathway from the Entorhinal Cortex Layer III (EC LIII; [36]), which are thought to carry, respectively, memorized and novel information [11, 8]. Classic theoretical models see phase precession, the correlation between the animal position and the firing phase of place cells within the theta cycle 28], as the generating mechanism of theta sequences [33]. This does not take into account the influence of dual input to CA1, which may act by itself as a considerable source of variability. It also neglects multiple results indicating how phase precession and sequences tend to appear most strongly under different network states, which in turn have been associated with prevalence of the EC LIII or CA3 inputs respectively 2, 38, 20]. While dynamical interactions between these inputs have been studied in detail [11, 2, 38, 15, 20], including via the use of activity perturbations [5, 30, 12, 39], dissecting their relative role is made difficult by the fact that EC LII also innervates CA3 through the tri-synaptic circuit. Thus, standard, unspecific manipulations of EC end up affecting both input streams [5, 30, 39, 25]. Here, we address this issue with a combination of ensemble physiology, layer-resolved oscillation recording and highly-specific manipulation of EC LIII projections to CA1.

We used the enhancer-driven gene expression (EDGE) [4] to create a novel transgenic mouse cross expressing an inhibitory opsin, JAWS [10], solely in EC LIII pyramidal cells (Figure 1a-h). Since EC LIII projects directly to the distal dendrites of CA1 pyramidal cells [36]), applying light to a region of CA inhibits the local direct input without affecting the circuit as a whole, or arguably even EC LIII cells themselves. We quantified expression levels by counting the number of JAWS-GFP RNA-positive nuclei within all brain areas and we estimated that the majority (78 %) were in EC LIII (Figure 1c). The expression of JAWS-GFP was also largely overlapping with Purkinje cell protein 4 (PCP4)-positive neurons (Figure 1d-h and Figure S1a-d), which can be used as a marker for EC LIII neurons. We recorded both large CA1 pyramidal cell populations and layer resolved oscillations with a novel version of the Hybrid Drive, combining 13 independently movable tetrodes, one linear silicon probe and one optic fiber in a single device ([19, 20], Figure 1i). Individual tetrodes were located in dorsal CA1 and the optic fiber was positioned above (*≈*100 µm) the pyramidal layer and enabled optical manipulation of the direct EC LIII projection in stratum lacunosum moleculare (s.l.m.) (Figure 1j,k,l,m and Figure S1e,f). Animals were trained, in a familiar environment, to run back and forth on a linear track collectng food rewards at both ends. The light of the laser was delivered continuously (max 30 s, 20 mW at fiber tip) every other lap, for both running directions consecutively (Figure 1n and Figure S2).

**Fig. 1.**
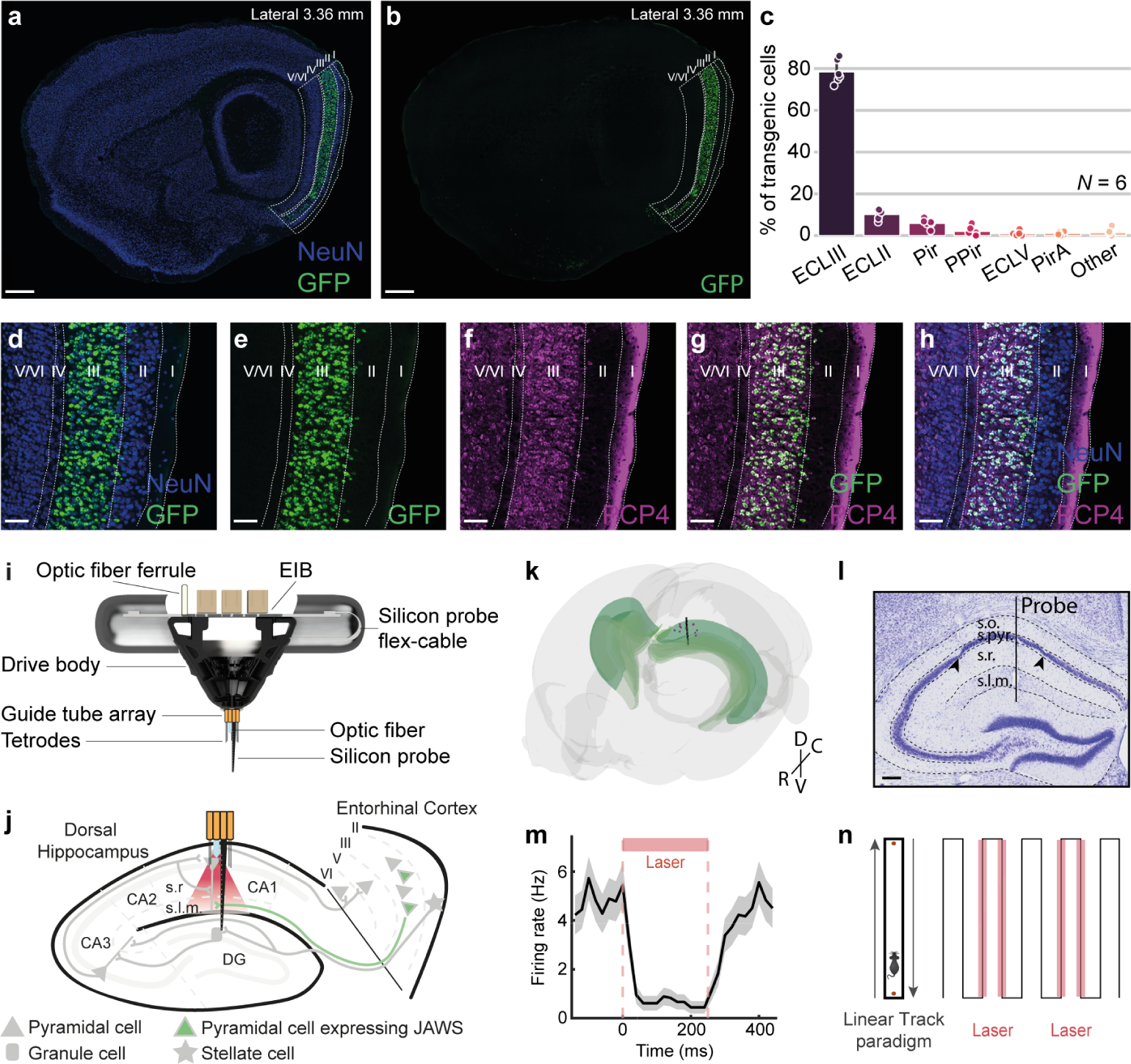
Novel transgenic cross for optogenetic manipulation of entorhinal cortex layer III and layer resolved ensemble electrophysiology in CA1. (a) Fluorescent in situ hybridization of a sagittal brain slice targeting the transgene GFP (green), fused to the JAWS opsin in the EC LIII-EDGE x JAWS mouse cross. Expression is largely restricted to Entorhinal Cortex Layer III (EC LIII). Counterstain NeuN (blue). Scale bar: 500 *µ*m. (b) Same as (a) without the NeuN counterstain. (c) Percentage of GFP labelled neurons by brain region. EC LIII, entorhinal cortex layer III; EC LII, entorhinal cortex layer II; Pir, piriform area; PPir, postpiriform transition area; EC LV, entorhinal cortex layer V; PirA, piriform-amygdala area. (d-h) High magnification images of dorsal MEC showing NeuN (blue), GFP (green) and PCP4 (violet) expression. Scale bar: 100 *µ*m. (i) Simplified schematic of the Hybrid Drive, with optic fiber. (j) Schematic of the hippocampus and entorhinal cortex showing tetrode bundle, silicon probe and optic fiber location in CA1 and specific expression of the JAWS opsin to a subset of EC LIII pyramidal cells projections. (k) 3D reconstruction from histology of tetrode (purple dots) and silicon probe (black shank) tracts. The hippocampal formation is depicted in green. D, dorsal; V, ventral; C, caudal; R, rostral. (l) Representative Nissl staining showing tetrode (black arrows) and silicon probe (balck bar) tracts from the Hybrid Drive implant. Scale bar: 100 *µ*m. s.o., *stratum oriens*; s.pyr., *stratum pyramidale*. (m) Representative example of suppressed firing in CA1 principal cell following repeated laser stimulation (250 ms). (n) Representative schematics of the linear track paradigm.

We first examined the effect of EC LIII distal input suppression on the rate code of CA1 neurons (Figure 2a). Among pyramidal neurons we only selected those with significant spatial modulation. Place fields were then selected and divided into two groups: “locking” (n=103) and “precessing” (n=199), on the basis of their relationship with theta oscillations (as previously described in [20]) during laser OFF periods. While we did not find changes in peak or mean (only a small, marginally significant effect for the precessing group), we found that the place field size slightly but significantly decreased during our manipulation (Figure 2b-d; t-test, p<0.001), only for the precessing sub-population. Effects were largely independent of changes in the running speed of the animals (Figure S3).

**Fig. 2.**
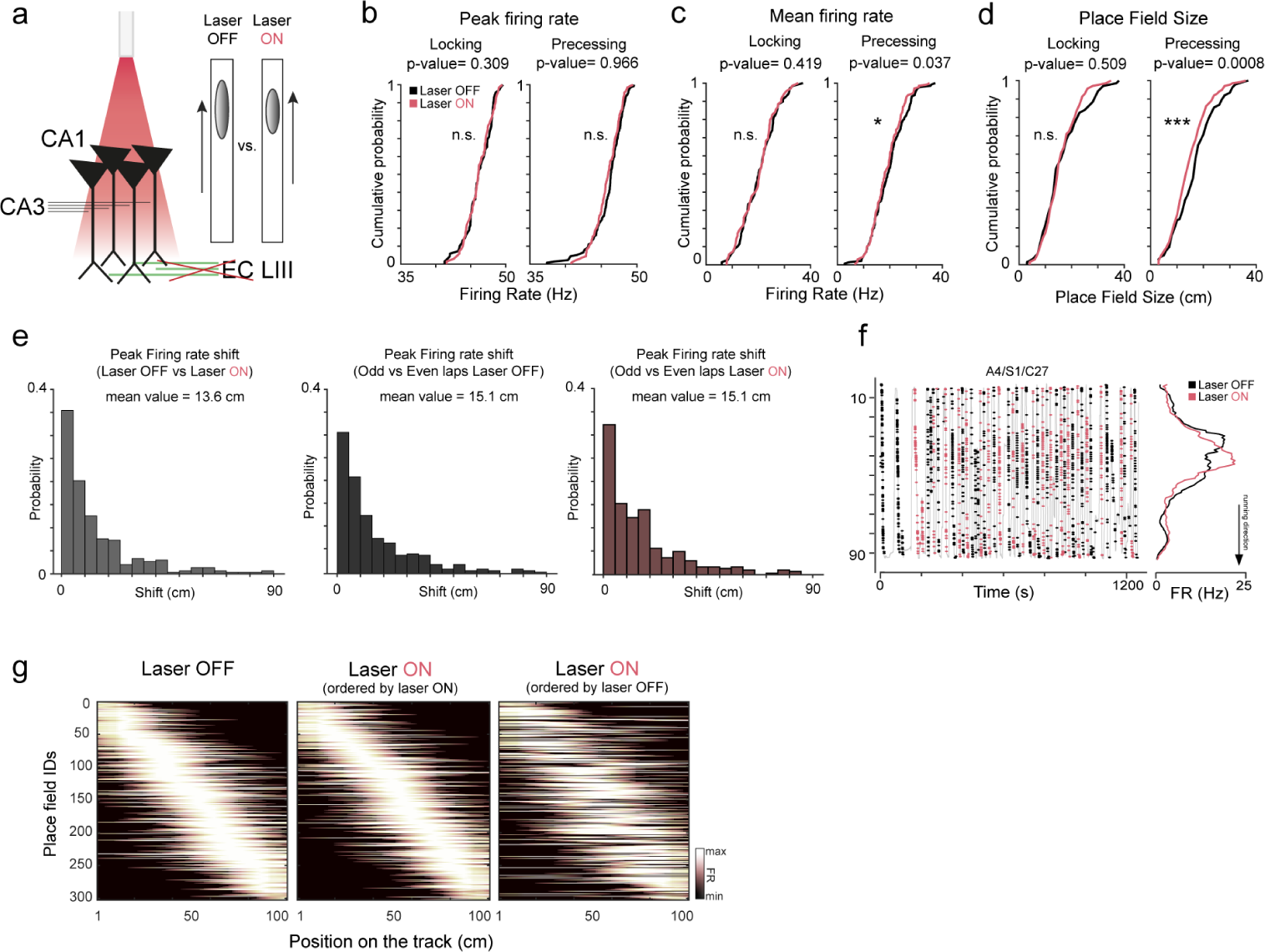
Spatial coding in CA1 does not rely on EC LIII direct input. (a) Schematic illustration of the optogenetic manipulation, selectively blocking the pre-synaptic terminals coming from EC LIII, on the distal dendrites of CA1 pyramidal cells in the s.l.m.. (b-d) Distributions of peak firng rate, mean firing rate and place field size in laser OFF (black) vs laser ON (red) conditions for phase locking and phase precessing fields. (e) Left, amount of shift of the place field center (defined as peak firing rate) between the laser OFF and laser ON conditions. Mean value = 13.6 cm. Center, change between Odd and Even laps in the laser OFF condition. Mean value = 15.1 cm. Right, change between Odd and Even laps in the Laser ON condition. Mean value = 15.1 cm. (f) Representative example of place field on the linear track. Grey: position of the mouse as a function of time, overaid with spiking activity only in the specified running direction (black dots: laser OFF. Red dots: aser ON). Right: average place field calculated from lap-by-lap spiking activity. (g) All place fields (rows), normalized by their maximum peak firing rate, sorted by their peak firing position on the inear track. *p*<*0.05, **p*<*0.01, ***p*<*0.001.

Importantly, place field representations remained largely stable, between the laser ON and OFF condition, as place field centers shifted by similar amount as in within-condition controls (Figure 2f and S5,). These results point to the fact that the EC LIII direct input is not essential for the consistent expression of a place field representation of a familiar environment (Figure 2g), and only a subset of place fields are affect by its inhibition.

We then asked how the inhibition of EC LIII inputs affects temporal coding processes within the CA1 network. This is likely to be state dependent, as the intensity of CA3 and EC inputs have been found to fluctuate in time. In fact, the magnitude of slow gamma (SG; 20-45 Hz) and medium gamma (MG; 60-90 Hz) oscillations correlate with the relative instantaneous strength of, respectively, CA3 and EC inputs in CA1 [11, 31, 15, 20]. We found that CA1 single cell temporal coding is perturbed by inhibition of EC LIII inputs (Figure 3a-d), lending interventional confirmation for this link. The suppression of phase precession appears to be due to the shift/suppression of spikes in the earliest phases of the theta cycle during periods of EC LIII inputs inhibition (Figure 3b and S6). It has been previously shown [2, 20] that theta phase precession is more strongly observed during elevated MG periods. Here we show that EC LIII inputs inhibition abolishes phase precession in precessing fields even during periods of spontaneously elevated MG network-states (Figure 3c-d). Spike densities of phase locking fields in the position-by-theta phase plane remain unaffected by our manipulation (Figure 3d). On a cell-by-cell basis, EC LIII inhibition removes any dependence of the phase-position slope on theta index [20] (Figure 3d, left panel), rendering phase precessing and locking fields indistinguishable in the phase plane at the single cell level. Together, these observations show how the EC LIII afferents are crucial for the normal expression of theta phase precession in CA1. Conversely, our manipulation did not cause any alteration in rate and temporal coding properties of place cells in control littermates, mice that did not express the inhibitory opsin but received the same light stimulation protocol (Figure S7).

**Fig. 3.**
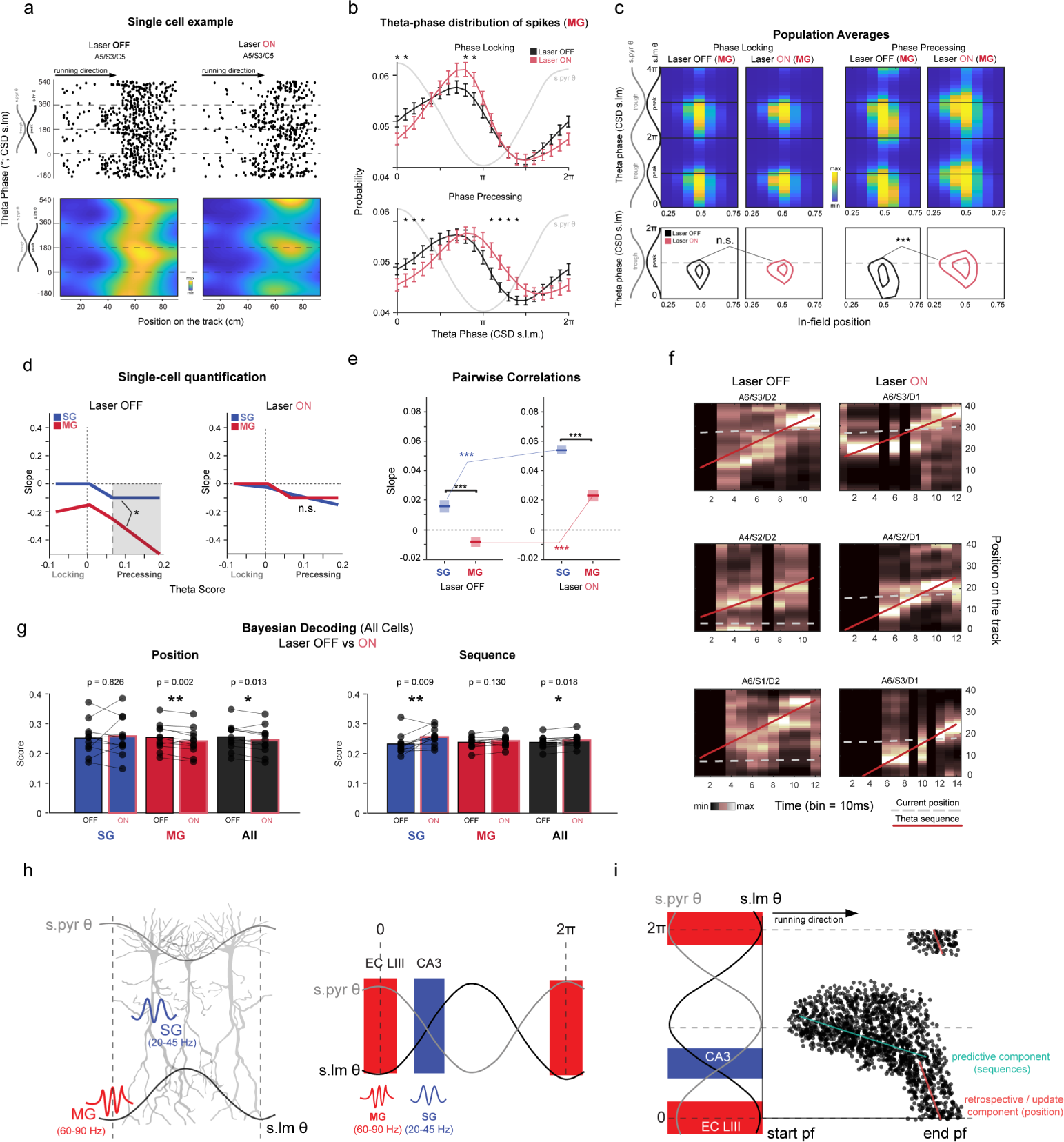
EC LIII input inhibition suppresses theta phase precession but enhances theta sequences. (a) Representative examples of impaired phase precession during periods of optogenetic manipulation (laser ON). Top, Theta phase–position raster plot, bottom phase-position density plot. 0 = peak of theta oscillation in the pyramidal layer. (b) Theta-phase firing probability distribution during MG-network states for phase locking and phase precessing populations, during laser OFF and laser ON periods (*p *<*0.05, t-test, Multiple Comparison Correction). (c) Top row, global average of the in-field spike probability during medium gamma network states, in Laser OFF (left) and Laser ON (right), for phase locking and phase precessing fields (two theta cycles). Phase 0 = peak of theta oscillation in the pyramidal layer. Bottom row, contour of regions with highest density of spikes for phase locking and phase precessing fields, curves correspond to 90 % and 85 % of the peak value of spike density (same data as top row panels). 2-Dimensional Kolmogorov–Smirnov test for spike densities (p*<*0.001). (d) Phase-position slope values plotted as a function of the Theta Score for each individual field, for periods of elevated SG and MG. Left, laser OFF. Right, laser ON. (p*<*0.05, Spearman correlation of Slow vs Gamma ratio vs Slope). (e) Place field distance vs phase distance regression for periods of elevated SG or MG and laser conditions (OFF and ON). All comparisons are statistically significant. Confidence intervals, p*<*0.001. (f) Examples of encoded position (y axis) reconstruction over a theta cycle (x axis, 1 bin = 1_1_0_9_ms). Black-white color scale: posterior probability from Bayesian decoding. Dotted grey line: real position of the animal, used to compute the amount of local coding as the probability density around the line. Red line: best positive-slope fit of probability distribution, used as an estimation of sequential non-local cell activation. (g) Comparison of mean positional and sequential scores from Bayesian Decoding between different gamma conditions (SG in blue, MG in red, all periods in black). Paired t-test. *p*<*0.05, **p*<*0.01. (h) Left, illustration summarizing the layer-resolved preferred phases for the CA1’s gamma components, and firing phase preference of CA3 and EC LIII afferent input [31]. (i) We redefine a dual-input model in which CA1’s phase precession can be conceived as an heterogeneous phenomenon that arises in a subset of place fields following the interaction between local CA1 firing dynamics and CA3/EC LIII input. We propose that the predictive component of precession ([24]), which consists of a sequence of spatial position with small variations in phase at the entrance and center of the place field, directly arises as a consequence of CA3-CA1 interactions (and more broadly through the tri-synaptic loop). Sequences that arise from these interactions are largely independent of the expression of strong phase precession pattern [26, 12], that instead rely on both CA3 and EC LIII input, occurring at earlier phases and toward the end of the place field (or procession [35], renamed here “retrospective/update component” [17, 35]).

Since its discovery, phase precession, and the resulting temporal distribution of place field spikes, has been proposed as the “engine” underlying the formation of theta sequences [33]. This would predict that EC LIII inputs inhibition, which disrupts phase precession, should also abolish theta sequences. We first tested this prediction by quantifying the correlation between field spacing and average theta phase separation of place cell pairs, an index of how consistently neurons fire in a stable sequence [13]. In contrast to the prediction above, the spatio-temporal correlation actually improved during periods of EC LIII inhibition (laser ON; Figure 3e), regardless of whether SG or MG dominated the LFP (p<10e-7, confidence intervals). Elevated SG-network states retained the highest spatio-temporal ordering in both conditions.

We confirmed and expanded these results by applying a Bayesian decoder (Figure 3f-g), looking at coding of current animal position and sequence expression across CA1 populations. Consistent with the pairwise result, we found an enhancement of sequential coding during periods of inhibition of EC LIII input (Figure 3g right panel; Paired t-test, p=0.018). In parallel, the accuracy of decoded position decreased, especially during MG network states (Figure 3g left panel; Paired t-test, p=0.002 and p=0.013). Additionally, sequential decoding improved in the second half of the theta cycle (Figure S9b, right panels) while positional decoding decreased in the first half of the theta cycle during our manipulation (Figure S9a left panels). This apparently surprisng result is in fact consistent with the partial dissociation between the expression of precession and sequences previously reported in interventional [26, 12] and descriptive 20] studies.

We have previously shown that precessing and locking populations form two groups of neurons that express theta sequences in a largely independent fashion [20]. Here, overall correlations increased during EC LIII inhibition, with cells from both groups expressing the highest ordering during slow gamma states (Figure 3d and Figure S8a). Phase precessing cells improved their spatio-temporal correlations with the phase lockng populations during medium gamma states (Figure S8a, right panel), while phase ocking cells improved their correlations both with cells of the other group and among themselves (i.e. with phase precessing cells; left panel). Notably, when EC LIII inhibition is applied, within and across group correlations became largely indistinguishable S8b). Thus, by releasing a subset of place fields from the influence of the EC LIII nput, sequences are expressed in a coherent way across the network, eliminating the differentiation between groups observed under normal input conditions.

Last, we explored the effects of the manipulation on global oscillatory rhythms. We did not find any change in the power profile in the s.l.m. (Figure S10). We found no alterations in the theta power profile (6-10 Hz) across all CA1 layers (pyramidal, radiatum and lacunosum moleculare; Figure S11). Similarly, theta-gamma phase-frequency coupling, was also left intact by our manipulation (Figure S12). These results suggest that our spatially restricted manipulation did not impair the global coordinated synaptic activity in CA1, beyond the small pool of neurons we are recording action potentials from.

Theta sequences are a fundamental organizing principle of hippocampal activity 6], which has been suggested to support planning and predictive coding [24]. Indeed, disrupting hippocampal sequences lead to impairment in planning behavior [25]. Our data suggests an update to current theories of sequence generation, decoupling them from theta phase precession. Phase precession has been seen as a unitary organizing principle for hippocampal neural activity, even generating a temporal phase code for animal position [22] more precise than the information contained in the firing rate. A number of potential network mechanisms have been proposed (for a review [14]), each with implications for hippocampal computations. We show instead that phase precession in CA1 is an heterogeneous phenomenon and may be understood as a consequence of the temporal distribution of two inputs, from CA3 and EC LIII (Figure 3h [9, 15]). Namely, EC LIII exerts tight control over the CA1 spiking activity in the earliest phases of the ongoing theta cycle and in the second part of the place field [23, 20]. Phase precession is best observed when spikes in both earlier and later theta phase, for which presumably the CA3 input is sufficient, are present. Thus, our data suggests that rather than a “phase code” for position spiking, theta phase precession is a consequence of the different distribution in space and theta phase of the CA3 and EC LIII inputs [9, 15]) (Figure 3h), with less immediate computational implications. Even more striking is the finding that EC LIII inhibition improves the organization of theta sequences. The most parsimonious interpretation of this result is that sequence generation is internal to the hippocampus proper, and probably due to interactions across the tri-sinaptic circuit (originating from EC LII). CA3 is endowed with the optimal recurrent connectivity to hold a reservoir of short-time spatial sequences, or preexisting neuronal dynamics [34], which may be passed to CA1. These spiking sequences may represent the basic units of many computations. Indeed, spikes in the first part of the place field, more robustly supported by CA3 input, appear to have a predictive coding character [24, 21]. CA3 input is also fundamental for rate and place coding in CA1 [26, 12, 39]. In contrast, the EC LIII input seems to be extraneous to such precise temporal organization, so that its presence disrupts sequence decoding. A hypothesis compatible with our data is that EC LIII drives plasticity in CA1 carrying external information, that may be used for prediction validation [17] and online update of CA1’s representations. This plasticity however cannot depend on precise spike timing, but it may rely on mechanisms at a longer time scales, as it has been recently proposed, bridging the *≈*100 ms gap between theta phases [3]. What is then, one may ask, the role of spreading CA3 and EC inputs at different phases of the theta cycle? It is possible that it is just the by-product of the feedback dynamics among the structures that produce theta oscillations (in the hippocampus and elsewhere) [7, 27]. On the other hand, we may speculate that such arrangement in the temporal segregation of inputs may provide a time window (the later theta phase), in each theta cycle, where sequences can be expressed “undisturbed”, favoring their readout in downstream structures. As it has been hypothesized [24, 21] the theta cycle may be separated in a phase for “encoding” (early theta phase) with retrospective properties [35] and a phase for “retrieval”, prediction and other computations. Our data suggest a clear role for direct EC LIII inputs in the encoding phase, suggesting a novel network architecture.

## Extended data

### Subjects

10 male mice were used in this study. 9 mice were implanted with a Hybrid Drive 19] and one with a 4-tetrode drive for representative examples of CA1 pyramidal unit inhibition with repeated stimulations (Figure 1m). All animals received the implant between 16 and 31 weeks of age. After surgical implantation, mice were individually housed on a 12-h light-dark cycle and tested during the dark period. Water and food were available *ad libitum*. All experiments were performed in accordance with the Norwegian Animal Welfare Act and the European Convention for the Protection of Vertebrate Animals used for Experimental and Other Scientific Purposes. Protocols were approved by the Norwegian Food and Safety Authority (FOTS ID 17898).

### Transgenic mouse lines

We crossed two transgenic mouse lines in this study. The ECLIII-tTA driver line expressing the tetracycline transactivator (tTA) within a subpopulation of cells confined to the Entorhinal Cortex Layer III (EC LIII) was made using the Enhancer Driven Gene Expression method with enhancer LEC13-8A, mapped to the Dok5 gene 4]. This ECLIII-tTA driver line was crossed to a tTA-dependent JAWS-tetO line generated as follows: transgene JAWS-GFP-ER2 (Addgene plasmid #65016) was cloned with restriction enzyme BamHI and NotI into an injection plasmid containing TRE tight 2 promoter (Clontech, now Takara Bio International). The injection plasmid was linearized by enzyme digestion to keep the relevant elements but remove the bacterial elements of the plasmids. Linearized DNA was run on a 1% agarose gel and isolated using a Zymoclean Gel DNA Recovery Kit (Zymo research, D4001). The resulting final injection construct thus consisted of a TREtight2 promoter followed by JAWS-GFP-ER2 sequences, WPRE element and SV40 polyA (TRE-Tight-2-tetO-JAWS-GFP-ER-WPRE-SV40). Fertilized oocytes from strain B6D2F1 x B6D2F1 were injected with the purified DNA. Both injection constructs were generated by the Kentros group at the Kavli Institute for Systems Neuroscience and sent for pronuclear injections at the Transgenic Mouse Facility at the University of Oregon, Eugene, OR USA. The control group (N=3) consisted of littermates that did not express the nhibitory opsin JAWS but received the same light stimulation protocol.

### Implant device and surgical procedures

The fabrication of the Hybrid Drive was done as described earlier [19]. This version of the Hybrid Drive is equipped with a linear silicon probe, 13 tetrodes and an optic fiber was incorporated in the design to carry out in-vivo optogenetic manipulations. The optic fiber was placed at the center of the array and glued in place at a depth of 1 mm from brain surface, sitting above the pyramidal layer *≈* 100 µm, in order to maximize the light coverage in the s.l.m. with minimal disruption of the cell layer. Mice were anaesthetized with isoflurane (5 % induction and 1-2 % maintenance). Prior to implantation, mice were given a subcutaneous injection of analgesics (Metacam, 5 mg/kg and Temgesic, 0.1 mg/kg), plus a local anaesthesia (Marcain, 1 mg/kg).

Anaesthetized mice were secured in a stereotaxic frame. The skull was exposed, cleaned and dried. 3 stainless steel M0.8 screws were used to secure the drive (1 ground screw in the frontal plate, 1 screw in the parietal plate opposite to the drive, 1 screw in the occipital plate). Using a 0.9 mm Burr drill, a craniotomy was made over the right cortex (top-left corner at AP: −1.20 mm; ML: 0.6 mm relative to bregma; bottom-right corner at AP: −2.30 mm; ML: 2.10 mm relative to bregma). The dura was removed and the array of the drive was slowly lowered into the brain with the silicon probe shaft and the optic fiber already adjusted at the final depth. To prevent cement from entering the guide tube array and the exposed brain, the craniotomy was filled with sterile Vaseline before lowering the array. The drive was cemented onto the skull using dental adhesive (Superbond C&B, Sun Medical, Japan) and tetrodes were individually lowered into the brain (5 turns - *≈* 900 µm). Mice were allowed to recover from surgery for at least seven days before experiments began.

### Neural and behavioral data collection

Animals were transported to the recording room from post-surgery day 3 and electrophysiological signals were studied during a rest session in the home cage. Each day, tetrodes were lowered individually in 45/60 µm steps (1/2 of a screw turn), until clear physiological markers for the CA1 pyramidal layer were discernible (sharp wave ripple complexes during sleep or theta during locomotion). The target location was usually reached in 10 days. Electrophysiological signals from the silicon probe contacts helped to refine the final position of the tetrodes.

Electrophysiological data were recorded with an Open Ephys acquisition board [32]. Signals were referenced to ground, filtered between 1 and 7500 Hz, multiplexed, and digitized at 30 kHz on the headstages (RHD2132, Intan Technologies, USA). Digital signals were transmitted over two custom 12-wire cables (CZ 1187, Cooner Wire, USA) that were counter-balanced with a pulley-system. Waveform extraction and automatic clustering were performed using Dataman (https://github.com/ wonkoderverstaendige/dataman). Clustered units were manually curated using the MClust toolbox. During all experiments, video data was recorded using a video camera (The Imaging Source, DMK 37BUX273) mounted above the linear track.

### Behavioral paradigm

Each behavioural session was preceded and followed by a 60-minute rest session in the animal’s home cage (“Pre sleep” and “Post sleep”) to ensure units’ stability. Mice were placed at one end of a 1-meter long track and trained to run to the other end, collecting a food reward (a piece of Weetos chocolate cereal). Following the animal’s consumption of the reward, a new one was placed at the opposite end. A lap was defined as an end-to-end run in which the animal’s body started at the first 10 cm of the track and finished at the other end without returning to its starting place. When the animals consistently carried out the task with a high number of laps, after approximately 10 days of consecutive training, the session with the optogenetic manipulation was carried out. Recordings typically lasted between 20 and 30 minutes. Animals were familiar with the recording room and the proximal and distal cues around the linear track. Mice were not food nor water deprived.

### Optogenetics

For optical silencing of EC LIII projections to s.l.m. of the hippocampal CA1 subregion, an optic fiber (core diameter: 100 µm, numerical aperture: 0.37, Doric Lenses) with a cone termination was incorporated in the Hybrid Drive and unilaterally mplanted in the right hemisphere at a depth of 1 mm. Optogenetic stimulation was performed by the delivery of continuous red light (640 nm, 20 mW at fiber tip) by a diode laser (Coherent Obis LX 1185054). Laser power was carefully curated to obtain sufficient irradiance in s.l.m. while avoiding excessive, detrimental effects. The onset and offset of light delivery were triggered as the animal finished eating the reward from one end and upon arrival at the other end of the linear track, respectively. Each lap with light stimulation was followed by a non-stimulated lap, providing an internal control for the effect of light stimulation. Optogenetic silencing sessions were performed once the mice proficiently performed on the linear track.

### Histology

Following the end of experiments, tetrodes were left untouched. Animals were euthanized using an overdose of pentobarbital, and perfused transcardially with RNase free 4 % paraformaldehyde in PBS. The brains were extracted and stored in RNase free 4 % paraformaldehyde for 24 hours before being transferred into RNase free 30 % sucrose for 2 days. For each animal, the left hemisphere was used to monitor transgene expression and the right hemisphere was used to verify the location of tetrodes, silicon probe and optic fiber. Both hemispheres were sectioned sagitally in 30 µm thick slices using a Cryostat (Fisher Scientific, Cryostar NX70). Sections were divided into 6 series and stored in a *−* 80 °C freezer. For the identification of recording sites and the position of the optic fiber, the tissue was stained with Cresyl violet. The slides were coverslipped and brightfield images were taken using a scanner (Zeiss Axio Scan.Z1, Germany) at a magnitude of 5x. Tracts were assigned to the Allen Mouse Brain Atlas and visualised in 3D using MeshView (https://www.nesys.uio.no/MeshView/). To validate transgene expression, the tissue was stained using both fluorescent in situ hybridization and immunohistochemistry. The sections were hybridized overnight at 62 °C with a FITC-riboprobe for GFP (2.5:1200; Roche, Cat. 11685619910). Next, the sections were washed with a tris-buffered saline with tween (TBST) solution and ncubated in a blocking solution (600 µl MABT, 200 µl sheep serum, 200 µl 10 % blocking reagent; Roche, Cat. 285 No. 11096176001) at room temperature for 4 hours. The blocking solution was replaced with anti-FITC-POD (1:1000) and the sections were left to incubate overnight at room temperature. The following day, the tissue was washed with a TBST solution before the fluorescein signal was developed using Tyr-FL (1:50, PerKinElmer kit) for 45 mins at room temperature, then washed in a TBST solution. The sections were then stained for PCP4 and NeuN with antibodies. The tissue was washed with 1X phosphate-buffered saline (PBS) (2 x 10 mins), placed in a permeabilisation buffer (1X PBS + 0.3 % Triton-X100, 10 mins) and preincubated in a blocking buffer (1x PBS + 0.3 % Triton X-100 + 3 % bovine serum, BSA, 1 hour). The sections were then incubated at room temperature for 48 hours with primary antibodies: Rabbit anti-PCP4 (1:300, Sigma, HPA005792) and Guinea Pig anti-NeuN (1:1000, Millipore, ABN90P) diluted in a solution of 1x PBS + 0.3 % Triton X-100 + 3 % BSA. Following primary antibody processing, sections were washed with 1X PBS + 0.3 % Triton X-100 + 3 % BSA (3 x 10 mins), then incubated at room temperature overnight with secondary antibodies: Goat anti-Rabbit Alexa Fluor 546 (1:400, Life technologies, A11010) and Goat anti-Guinea pig Alexa Fluor 647 (1:400, Life technologies, A21450) diluted in a solution of 1X PBS + 0.1 % Triton + 1 % BSA. The slides were coverslipped and images were taken using a scanner (Zeiss Axio Scan.Z1, Germany) at a magnitude of 5x.

### Automated cell counting

Cell counting was performed on one series of brain slices per animal. The EBRAINS semi-automated workflow QUINT ([37, 18, 29]; https://ebrains.eu/service/quint/) was used for brain-wide cell quantification. The brain slices were registered to the Allen Mouse Brain Atlas (http://www.brain-map.org/) using the QuickNII software. Anatomical boundaries from the atlas were fine-tuned to fit the data using non-linear adjustments in VisuAlign (https://www.nitrc.org/projects/visualign/). Cells were extracted by using machine-learning image segmentation toolkit ilastik (inter-active machine learning for (bio)image analysis, https://www.ilastik.org/). Finally, Nutil Quantifier was used to quantify and assign anatomical locations to labelled cells obtained by segmentation of brain images.

### Place Cell Identification

We first selected putative excitatory pyramidal cells using their auto-correlograms, firing rates and waveform information. Specifically, pyramidal cells were classified as such if they had a mean firing rate <8 Hz and the average first moment of the auto-correlogram (i.e., the mean value) occurring before 8 milliseconds. Only cells classified as pyramidal cells were used for further place cell analysis. Place cells were defined applying a combination of different criteria. All analysis were performed on speed-filtered activity, after removing periods in which the animal speed was smaller than 3 cm/s. Only cells with an average activity above 0.3 Hz were taken into account. Then for each of these cells, the Skaggs information per second 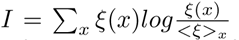 was compared to the distribution of information values resulting from randomly shuffling cell spike times. A cell passed the selection if its Skaggs information was significantly higher than the ones of the surrogate distribution. Last, only place cells with peak firing rate higher than 1 Hz were kept for further analysis.

### Place Cell Classification

Spikes were filtered both in speed (>3 cm/s) and in position, by taking only those emitted within the boundaries of the cell place field. For each place field, a ‘Theta Score’ [20] was calculated as follows: Theta Score = Precession Score - Locking Score. For the Precession Score we used a circular-linear correlation to measure the degree of dependency between the spike phase and the position within the field. For the Locking Score we computed the length of the Reyleigh vector associated to the distribution of spike phases with respect to the instantaneous theta phase. Each measure produced a score between 0 and 1, with 1 denoting perfect phase-position relation or perfect phase locking. Negative Theta Score values indicate that spikes probability are mostly modulated by specific phases of theta (‘phase locking cells’), while positive Theta Scores suggest a strong phase precession (‘phase precessing cells’).

### Place Cell Properties

Place cell properties were calculated using a Generalized Linear Model (GLM; described in detail in [20]) to infer the spike probability of each place field given i) the position within the place field; ii) the instantaneous theta phase and iii) the instantaneous power in a specific gamma range and layer. In all our analysis, we considered a partition of the environment in bins of 2.5 cm. Place fields were isolated as continuous regions in the rate map with rate higher than 20 % of the cell maximum activity. Spatial mean firing rate was calculated integrating these probabilities over the other variables. Spatial firing rate of each cell was calculated within each condition and separately for laser OFF and laser ON. As place field center we took the bin within the place field with highest firing rate. The effect of speed on the firing probabilities was addressed in a similar manner, by including the instantaneous velocity of the animal as a further covariate in the GLM. Place coding analysis were restricted to place field without substantial change in peak firing rate between conditions (<15 cm, based on average Peak Firing rate shift).

### Neural data analysis on oscillations

Before applying other analysis, continuous (LFP) signals were down-sampled to 1 kHz. Power spectral density (PSD) estimates were calculated across the frequency range 30–100 Hz using the pwelch function in Matlab, for each condition (laser OFF vs. aser ON) separately. LFP across the probe’s 16 contacts was also used to compute the corresponding Current Source Density (CSD) signal. This transformation is based on computing the discrete laplacian along the probe axis (that is along the direction running through CA1 layers). The resulting CSD signal is limited to 14 channels (16 minus the 2 extremes) CSD signals were used to compute the phase-amplitude coupling strength between a reference theta oscillation and faster oscillations, in a [15–150] Hz range. We first applied a wavelet transform to obtain the analytical signal over time and then we tracked the evolution of the phase-amplitude couplings across different ayers and frequencies, as previously described [20]. For each of the identified main gamma frequency ranges (slow [20–45] Hz and medium [60–90] Hz) we extracted layer-specific coupling strength and phase range of frequencies. Theta oscillation reference was generally taken from the s.l.m., where theta modulation of the LFP is strongest. Additionally, CSD signal were used to estimate the layer-resolved Gamma Coefficient as follows: the instantaneous balance between the power in the slow gamma (P*_Slow_*(t)) and medium gamma (P*_Med_*(t)) frequency range was computed as follow. First, both the slow gamma power and medium gamma power during running periods were separately z-scored. Second, a power-ratio score, spanning the [-1 1] interval was computed as:

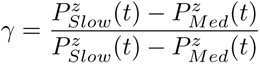

so that a value of −1 would correspond to total medium gamma domination and +1 to slow gamma completely dominating.

### Generalized-Linear Model (GLM)

The GLM analysis was carried out (separately for periods with and without laser manipulation) as previously described in [20].

### Pairwise Spike Timing correlations

We performed pairwise correlations analysis as previously described in [20]. Firstly, the analysis was done separately for periods with and without laser manipulation. For each cell, only the first spike in each theta cycle and only spikes emitted within the main place field (where the peak firing rate occurs), were considered. For each of these spikes the simultaneous slow/medium gamma power ratio was calculated, and then used as a label to further subdivide the spikes. Then, all the cell pairs in the population of simultaneously recorded place cells were considered. Each pair (A,B) was defined by 1) the spatial distance between the place fields of cell A and B, computed as the distance between the centers of mass (using the distance between the fields peaks did not change the results) and 2) the average phase interval between the spikes selected for the two cells. The latter was computed by taking spikes of cell A as reference and within each theta cycle (for which a spike from both cells was available) taking the dθ*_A−>B_*(k) (where k stands for the kth theta cycle), that is, the signed difference between the phase of the spike of cell A and that of cell B spike. The center of mass of the dθ*_A−>B_* distribution was then computed and used as a measure of the average phase offset between that cell pair. The spatial distance and phase distance were then organized in a two-dimensional space, and the average phase offset was calculated for a range of spatial distances, provided that there were at least ten pairs of cells available in that range. The analysis was repeated using cell pairs from different subgroups, and selecting spikes released under a particular gamma balance.

### Bayesian Decoding

Bayesian decoding analysis were carried out as previously described in [20]. First we separated periods with and without laser manipulation and analysed them independently from each other. We selected periods of active locomotion and segmented time using the peaks of theta oscillation (extracted from the stratum lacunosum moleculare). A sliding window of 30 ms length, with an offset of 10 ms, was used to further divide each theta period. For each of these sub-windows, we built a population vector 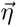(t), where t indicates the time within the theta cycle. Using Bayes’ formula, we computed P(x*|*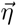(t)): the likelihood of position on the track x given the population activity.

The set of probabilities P(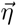(t))*|*x) were obtained from marginalizing the result of the GLM over the phase of spiking. The resulting probability density P(x, t) was normal-zed so that ΣP(x, t) = 1 for each time window t that contained 3 spikes or more.

Time windows with 3 or less spikes were excluded from further analysis. Similarly, theta cycles with less than n ‘active’ time windows were removed from the following analysis (n was set to 5 when considering the entire cell population and decreased to 3 when decoding from subgroups of cells). For each theta cycle we computed two scores, using the normalized probability density. The amount of ‘local’ information in the activity was first quantified by summing the probability concentrated around the current position of the animal ϒ*_Local_* = ΣP(x*^∗^*, t). x*^∗^* denotes the set of spa-*_t_*tial bins in a range of 5 cm of the animal position at time t. Running a set of linear regression over a modified version of the P *^∗^*(x, t) matrix, where the ‘local’ decodng probability density had been subtracted out of the original P(x, t) so that we could factor out the effect of positional decoding from the reconstructed trajectory), allowed us to detect the presence of ‘non-local’ sequence-like activity over a signifcant spatial interval around the animal position. The best-fit was identified among these regressions, overlapping with the largest amount of probability density. That is, ϒ*_NonLocal_* = max*_β_* Σ*_k_* P *^∗^*(x*_β_*_(_*_k_*_)_, t*_β_*_(_*_k_*_)_) where β indicates a set of linear parametrizations of x and t: x = β_1_ + t *×* β_2_. Importantly β_2_ was taken to be always > 0 so to exclude fits very close to the ϒ*_Local_* defined above. The same procedure was applied using different set of cells to compute the probability density, so to obtain an estimation of the spatial information carried by specific cell groups (the entire population and either phase precessing or phase locking cells). Theta cycles were further classified according to the simultaneously expressed gamma power ratio. To compare the nature of activity across different cell populations, the decoding scores obtained from each cell group were first normalized to max_Θ_ϒ = 1 (where the max is taken over all available theta cycles); then, for each theta cycle in which enough spikes were emitted by cells from both independent cell groups, we computed the difference between the two obtained decoding scores ϒ_1_ and ϒ_2_ (either Local or Non-Local) as the distance of the point (ϒ_1_,ϒ_2_) from the equal-score diagonal ϒ_1_ = ϒ_2_.

### Statistics

Data analyses were performed using custom MATLAB scripts (The Math Works). Paired and unpaired t-tests were performed using standard built-in MATLAB functions. 2-D Kolmogorov-Smirnov test was performed using custom code based on the Peacock algorithm. All tests were two-tailed, except where stated otherwise.

## Acknowledgments

We thank Morgane Audrain for help with the 3D illustration of the Hybrid Drive, Jordan Carpenter, Abraham Z. Vollan, the Kentros and Battaglia labs for comments on the manuscript and helpful discussions at different stages of the project. This work was supported by the European Union’s Horizon 2020 research and innovation program (MGate, grant agreement no. 765549; M.G., O.C., C.K. and F.P.B.), the European Research Council (ERC) Advanced Grant “REPLAY-DMN” (grant agreement no. 833964; F.P.B.), and European Union’s Horizon 2020 Research and Innovation Program Grant “BrownianReactivation” (grant agreement no. 840704; F.S.), FRIPRO ToppForsk grant Enhanced Transgenics (90096000) of the Research Council of Norway (O.C., Q.Z. and C.K), the Kavli Foundation (O.C., Q.Z. and C.K), the Centre of Excellence scheme of the Research Council of Norway—Centre for Biology of Memory and Centre for Neural Computation (O.C., Q.Z. and C.K), The Egil and Pauline Braathen and Fred Kavli Centre for Cortical Microcircuits (O.C., Q.Z. and C.K), and the National Infrastructure scheme of the Research Council of Norway—NORBRAIN (O.C., Q.Z. and C.K).

## Author contributions

Conceptualization: M.G., O.C., F.S., C.K. and F.P.B.; Investigation and Data Curation: M.G.and O.C.; Methodology and Software: M.G, O.C., Q.Z., F.S., C.K. and F.P.B.; Resources: C.K. and F.P.B.; Formal Analysis and Visualization, F.S., M.G. and O.C.; Writing – Original Draft: M.G., O.C. and F.P.B.; Writing – Review & Editing: M.G., O.C., F.S., C.K. and F.P.B.; Supervision and Funding Acquisition: F.S., C.K. and F.P.B.

## Ethics declarations

The authors declare no competing interests.

---

**Fig. S1.**
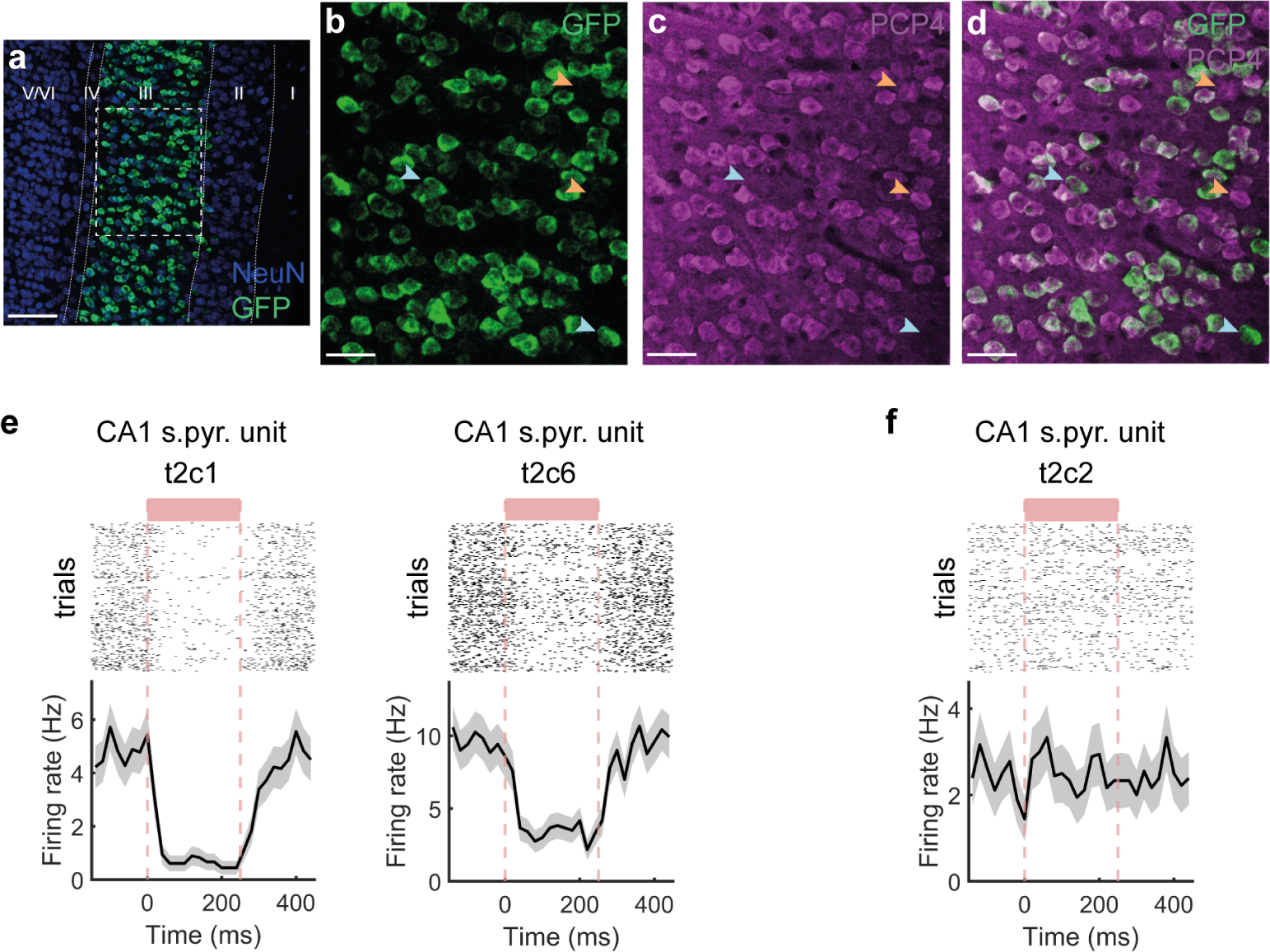
Expression of the JAWS opsin is largely specific to EC LIII excitatory neurons. (a) Fluorescent In Situ Hybridization targeting the transgene GFP and NeuN. Scale bar: 100 *µ*m. (b-d) Large majority of GFP/opsin expressing cells overlap with PCP4+ cells, marker for EC LIII neurons. Light blue arrows: example GFP+/PCP4-cells. Orange arrows: GFP-/PCP4+ cells. Scale bar: 50 *µ*m. (e) Additional representative examples of suppressed firing in CA1 principal cell following repeated laser stimulation (250 ms). Raster plot (top) and peri-stimulus time histogram (PSTH, below). Left plot, same unit as in Figure 1m, with individual manipulations trials included. (f) Same as in (e), but for CA1 principal cell not affected by repeated laser stimulation (250 ms).

**Fig. S2.**
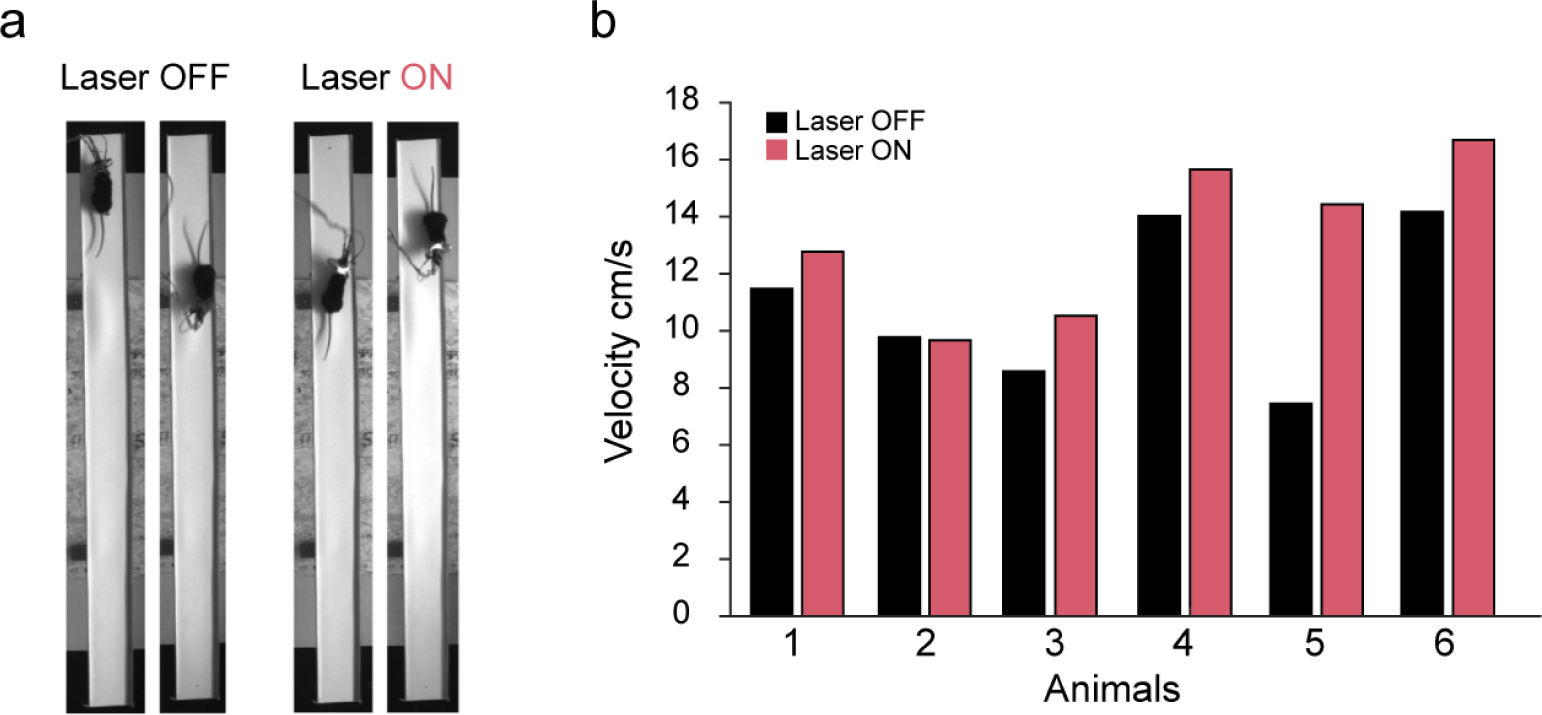
Velocity performances during the goal-directed linear track task. (a) Examples of mice implanted with a Hybrid Drive in the linear track task during either laser OFF or laser ON runs, for both running directions. (b) Barplots displaying velocity performances across conditions (average across sessions). Periods of immobility (movements *<* 3 cm/s) were removed.

**Fig. S3.**
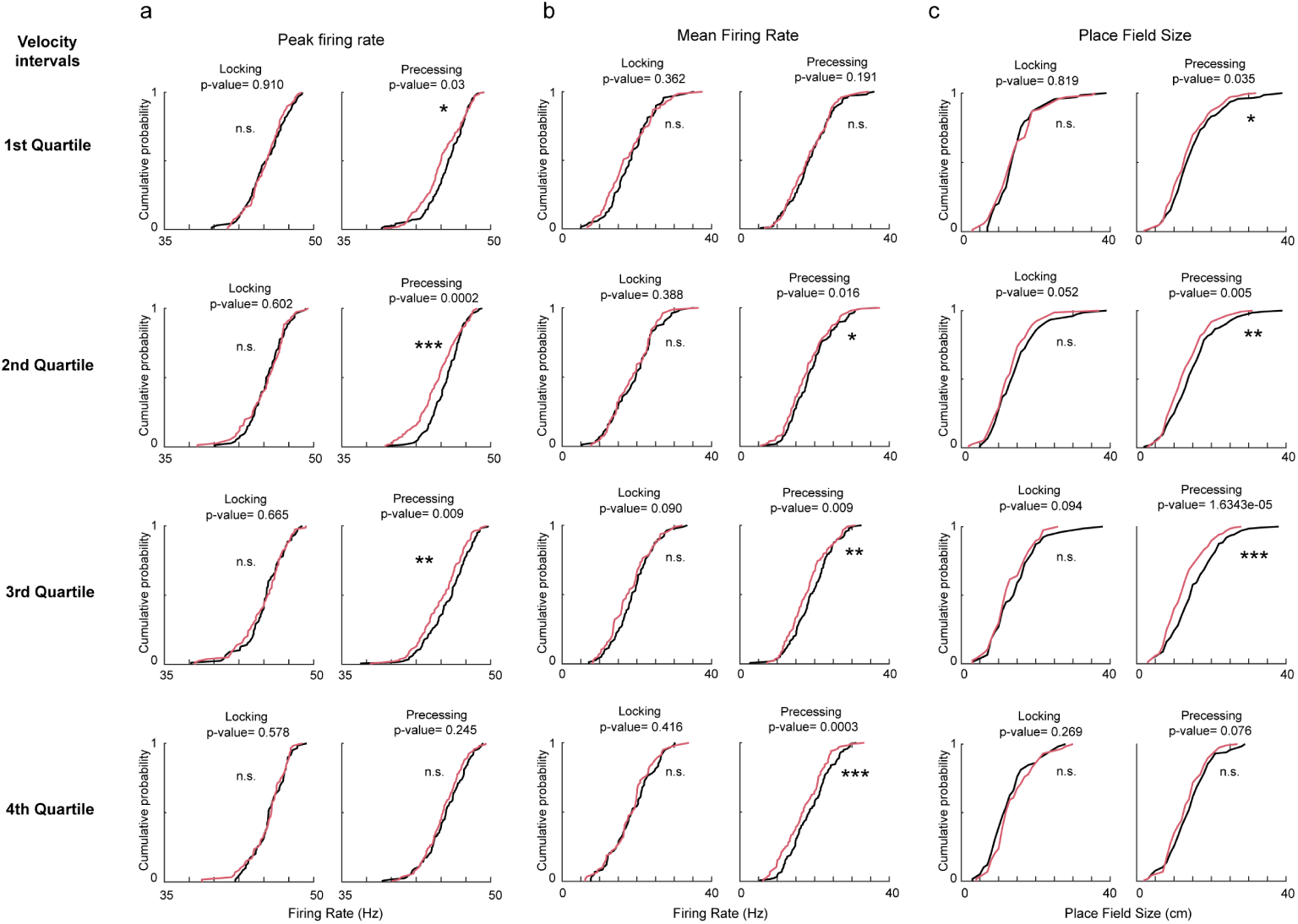
Rate and place coding properties across velocity intervals. (a-c) Distributions of peak firing rate, mean firing rate and place field size respectively, across different velocity intervals. Same amount of data in each velocity interval. *p*<*0.05, **p*<*0.01, ***p*<*0.001.

**Fig. S4.**
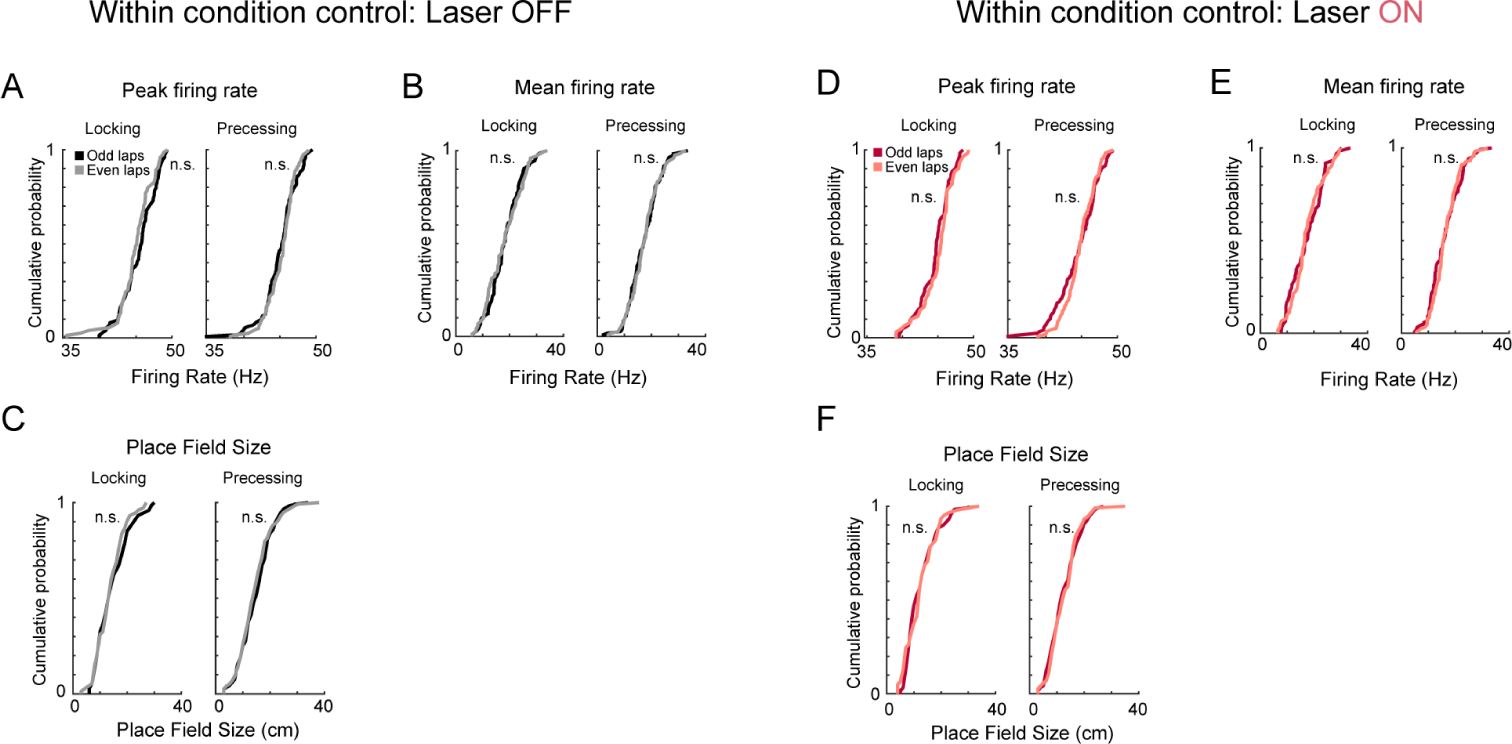
Rate and place coding properties within conditions. (a-c) Distributions of normalized peak firing rate, mean firing rate and place field size, respectively; using only within conditions data (Odd and Even laps for laser OFF and laser ON separately). No significant differences were found (p*>*0.05; t-test for all conditions all comparisons). (d-f) Same thing but using Odd and Even laps within the Laser ON condition. No significant differences were found (p*>*0.05; t-test for all conditions all comparisons).

**Fig. S5.**
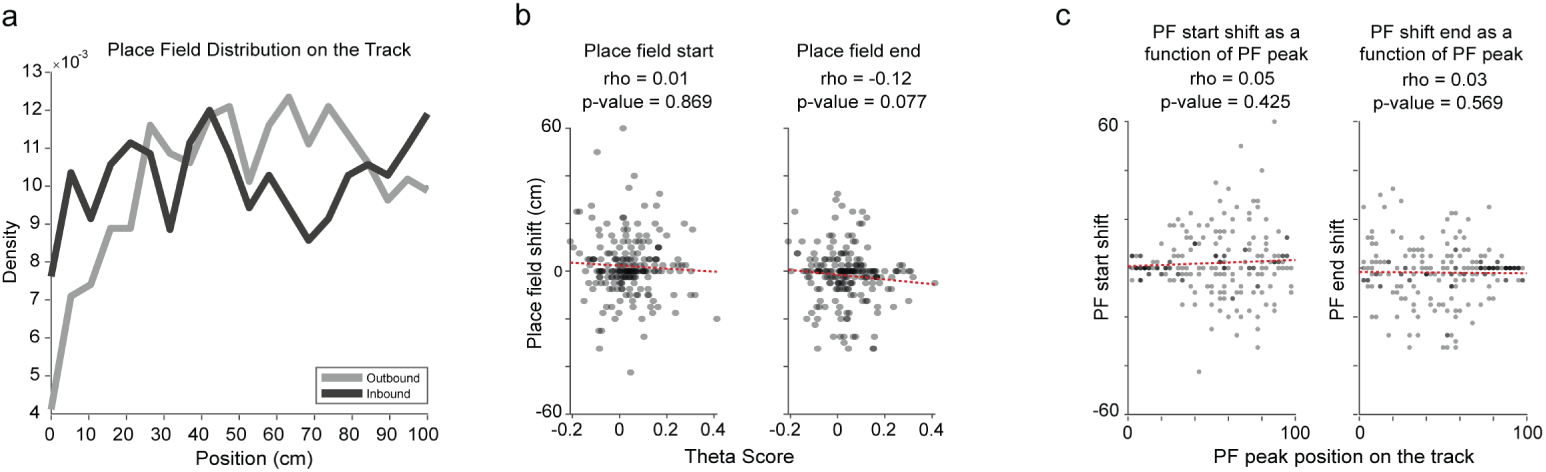
Place field properties on the linear track. (a) Distribution of place fields on the linear track, for both running directions separately. (b) Correlation between the place field shift (in cm) and the Theta Score, for the place field start and end separately (Spearman’s rho, p*>*0.05). (c) Correlation between the place field shift and the position of the peak firing rate on the linear track, for the place field start and end separately (Spearman’s rho, p*>*0.05).

**Fig. S6.**
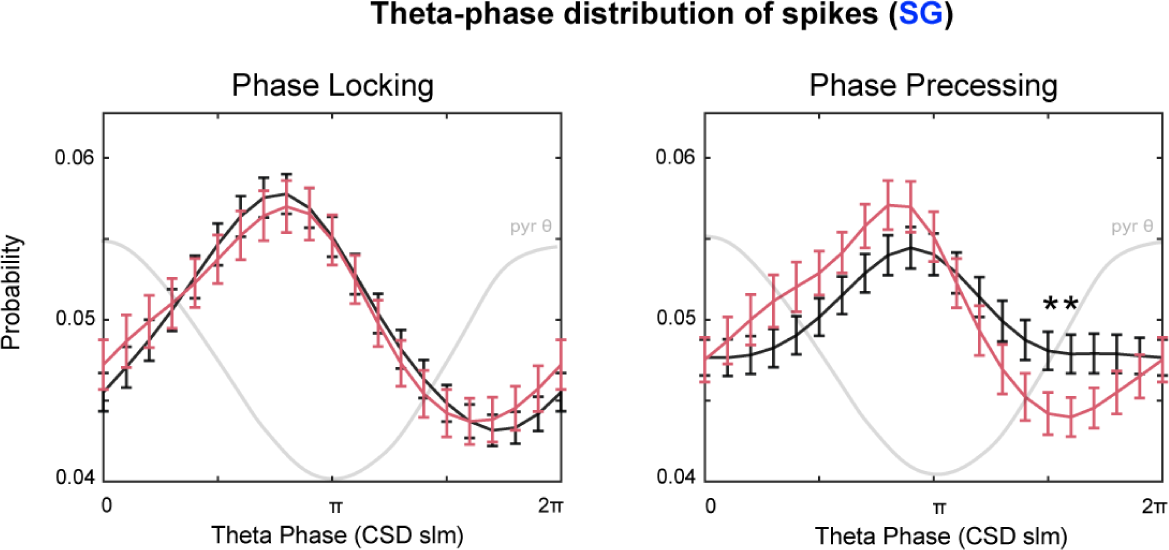
Theta-phase distribution of spikes during SG network states. (a) Theta-phase firing probability distribution during SG-network states for phase locking and phase precessing populations, during laser OFF and laser ON periods (p*>*0.05, t-test)

**Fig. S7.**
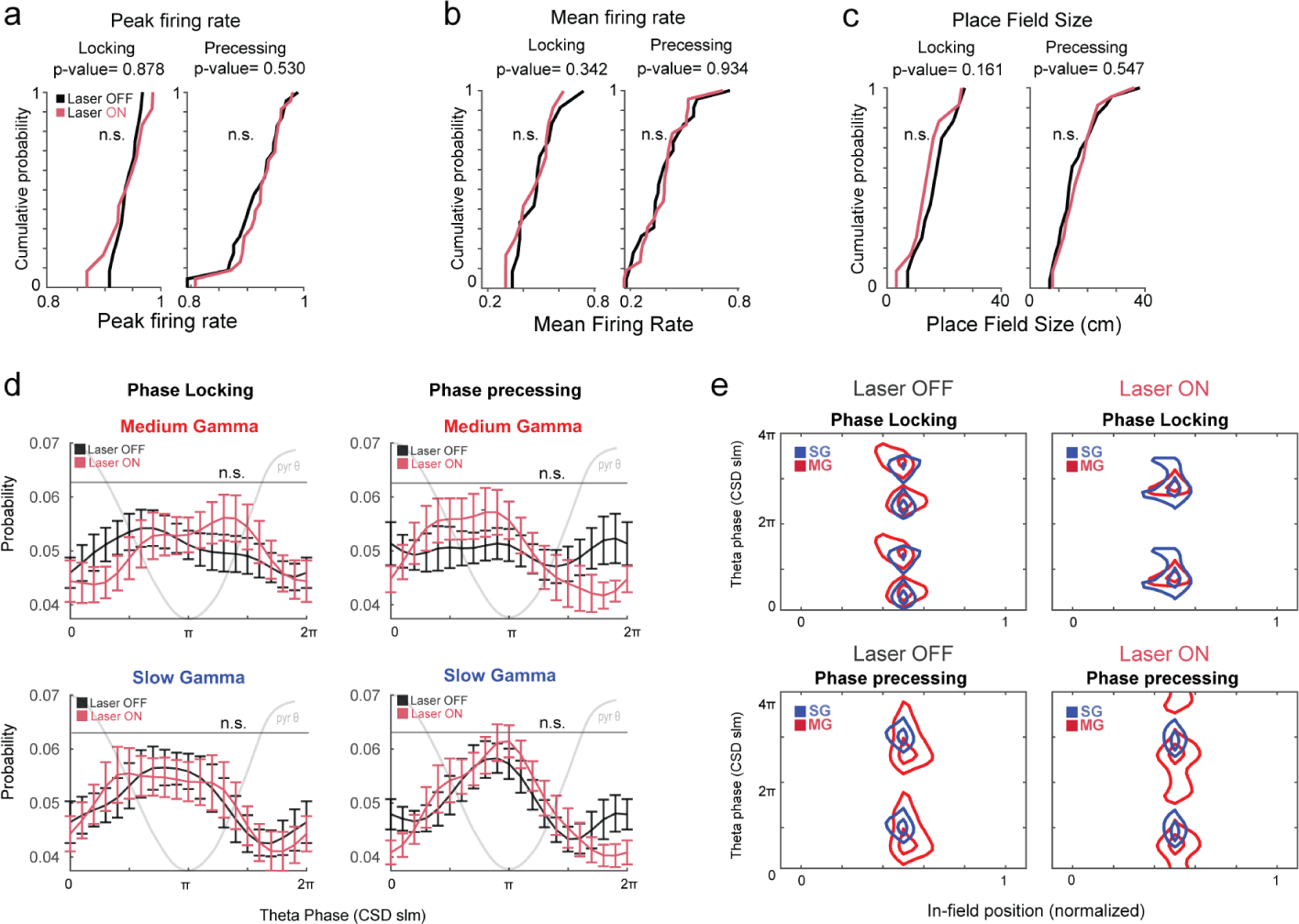
Rate and phase code dynamics in single positive control animals. (a-c) Distributions of normalized peak firing rate, mean firing rate and place field size in laser OFF vs laser ON conditions. All comparison are not statistically significant (p*>*0.05; t-test for all conditions, all comparison). (d) Theta-phase firing probability distribution during MG-and SG-network states for phase locking and phase precessing populations, during laser OFF and laser ON periods (p*>*0.05, ttest, Multiple Comparison Correction). Animals (N=2); session (n=5). (e) Global average of the in-field spike probability during medium gamma network states, in laser OFF (left) and laser ON (right), for phase locking and phase precessing fields. Phase 0 = peak of theta oscillation in the pyramidal layer. Contour of regions with highest density of spikes for phase locking and phase precessing fields. Curves correspond to 90% and 85% of the peak value of spike density. 2-Dimensional Kolmogorov–Smirnov test for spike densities (p*>*0.05). Animals (N=2); session (n=5).

**Fig. S8.**
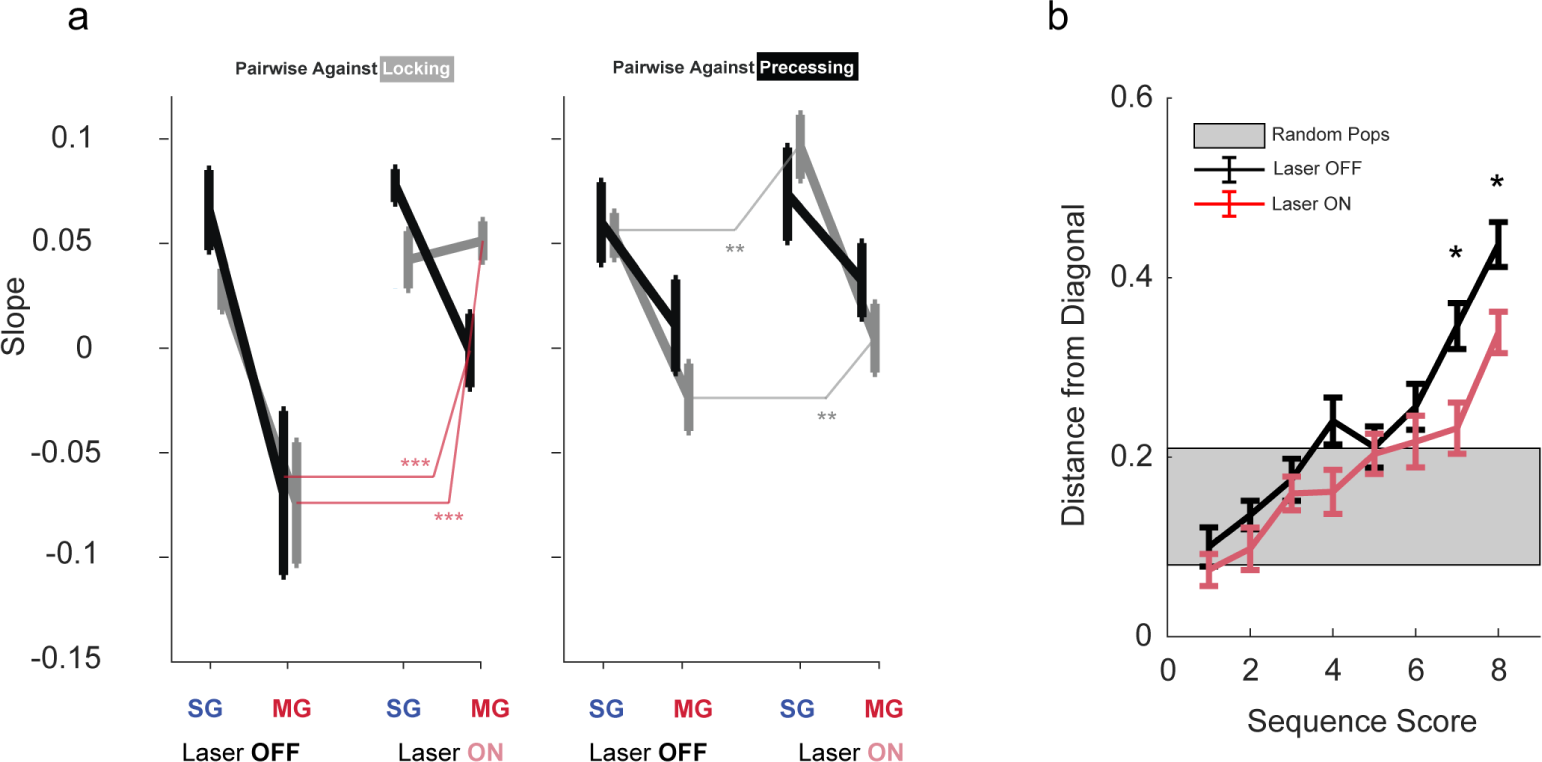
Pairwise correlation across place cell groups. (a) Spatio-temporal pairwise spike correlation computed for cell pairs either within the same cell group or across groups, as a function of the instantaneous gamma balance (SG and MG) and the laser conditions (OFF and ON). (b) Distribution of simultaneously measured sequence decoding scores (left: position; right: sequence) rom locking and precessing population activity on a theta cycle by theta cycle basis. Equivalent decoding scores correspond to value of 0 on the y-axis, while increasing values indicate a larger discrepancy of the decoded sequence quality in the two populations. Decoding events are arranged on the x-axis according to their average quality, and average is performed over each bin. Shaded area is estimated from discrepancy distribution obtained from randomly reassigning locking and precessing cells in new sub-populations (t-test p=0.01, laser Off vs laser On). Data are presented as mean ± SEM.

**Fig. S9.**
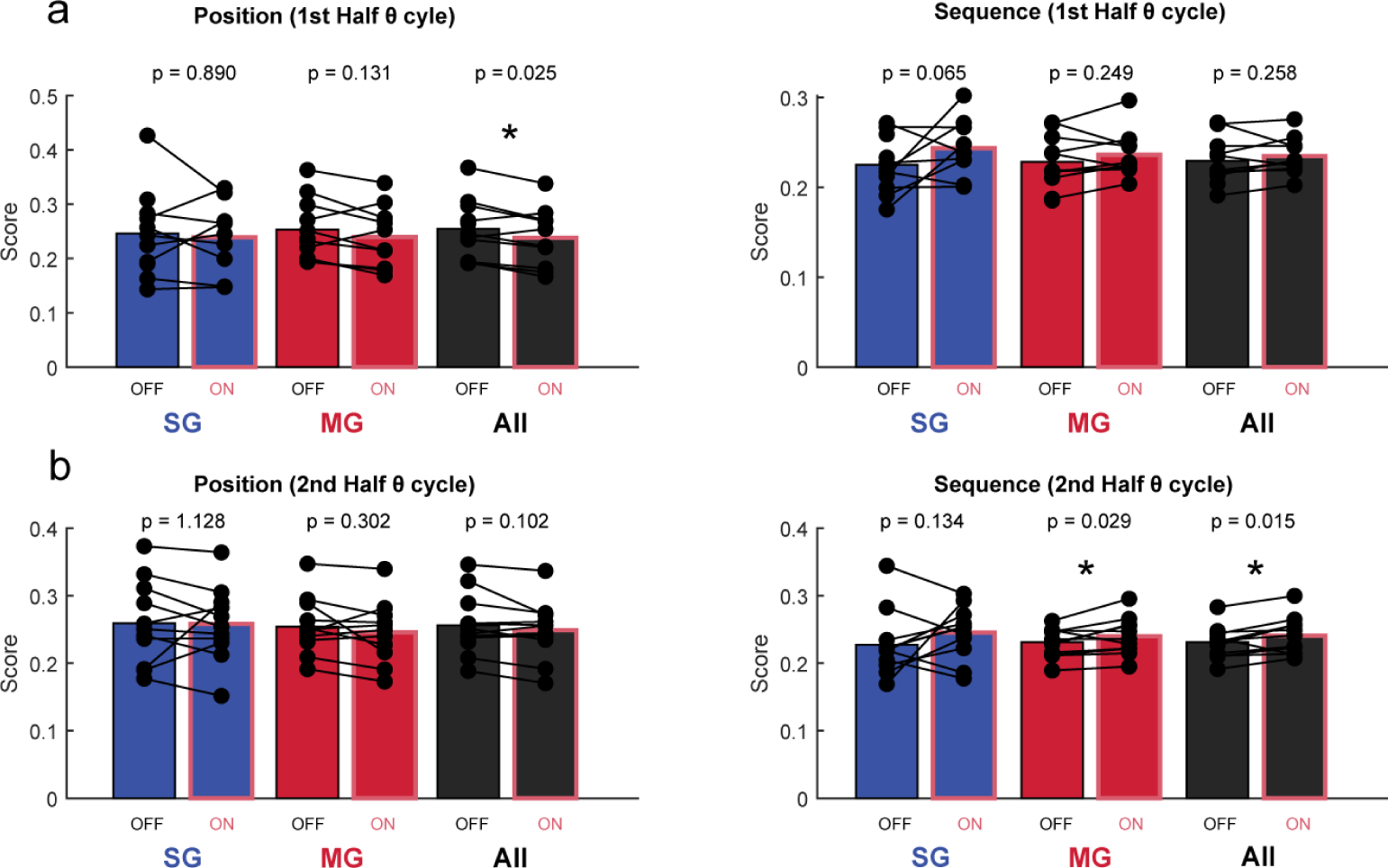
Bayesian decoding across the theta cycle. (a) Comparison of mean positional and sequential scores between laser OFF vs laser ON periods in different gamma conditions (slow gamma in blue, medium gamma in red, all periods in black) in the first half of the theta cycle. Paired t-test. Statistical values in order of appearance, *p*<*0.05. Position: p=0.890, p=0.131, p=0.025. Sequence: p=0.065, p=0.249, p=0.258. (b) Comparison of mean positional and sequential scores between different gamma conditions (slow gamma in blue, medium gamma in red, all periods in black) in the second half of the theta cycle. Paired t-test. Statistical values in order of appearance; **p*<*0.01, *p*<*0.05. Position: p=1.128, p=0.302, p=0.102. Sequence: p=0.134, p=0.029, p=0.015.

**Fig. S10.**
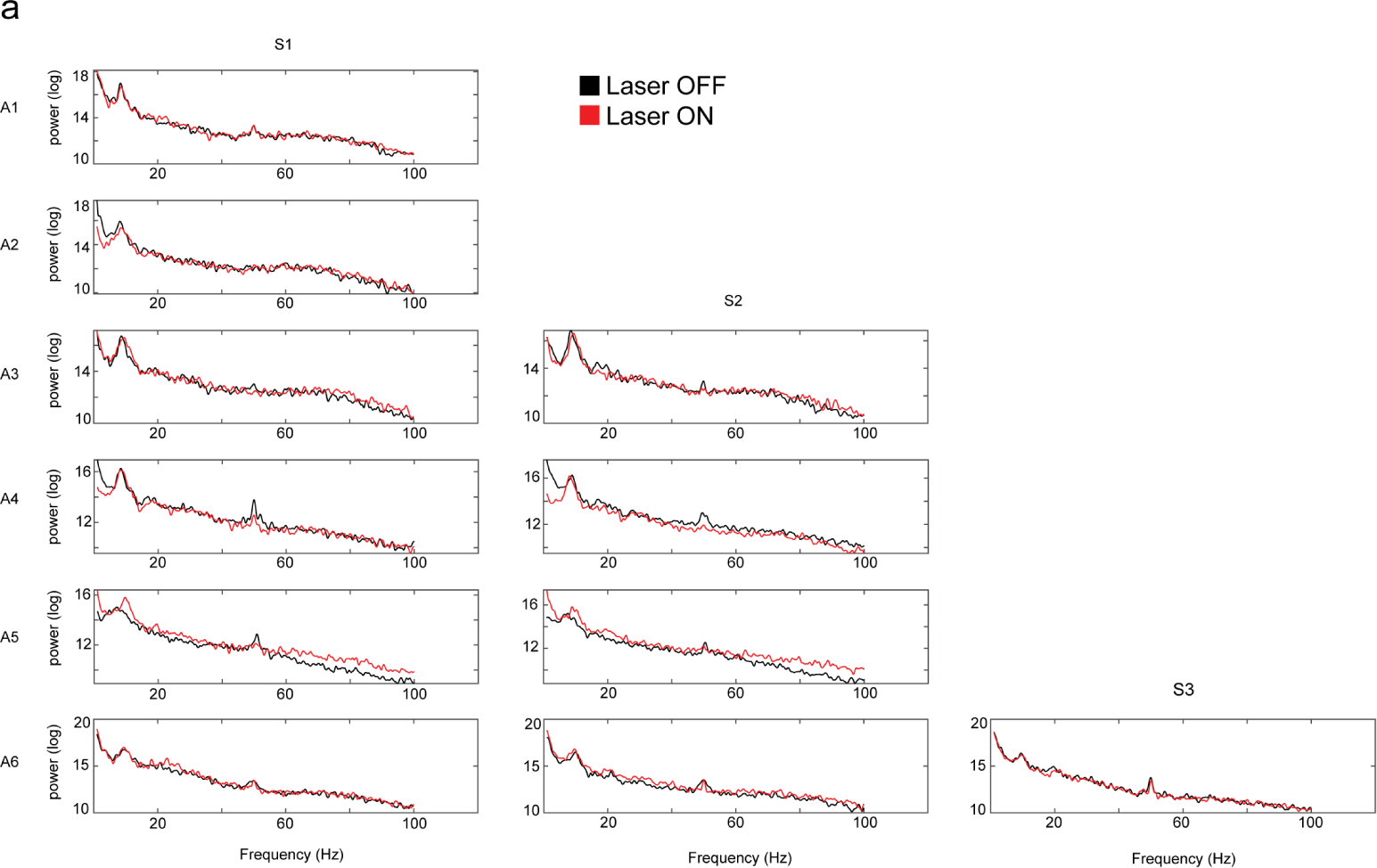
Power spectral density estimate in CA1 during optogenetic inhibition of EC LIII direct input. (a-b) Power spectral density estimate of LFP signals in s.l.m.. No differences are ound between conditions (laser OFF vs laser ON). Periods of immobility (movements *<* 3 cm/s) are removed. All animals (N=6), all session (n=11). Higher power profiles in both sessions of the Animal 5 are due to higher velocity performances during laser OFF and laser ON runs (Figure S2) - faster velocities coincide with higher power in hippocampal oscillations at all frequency bands [1].

**Fig. S11.**
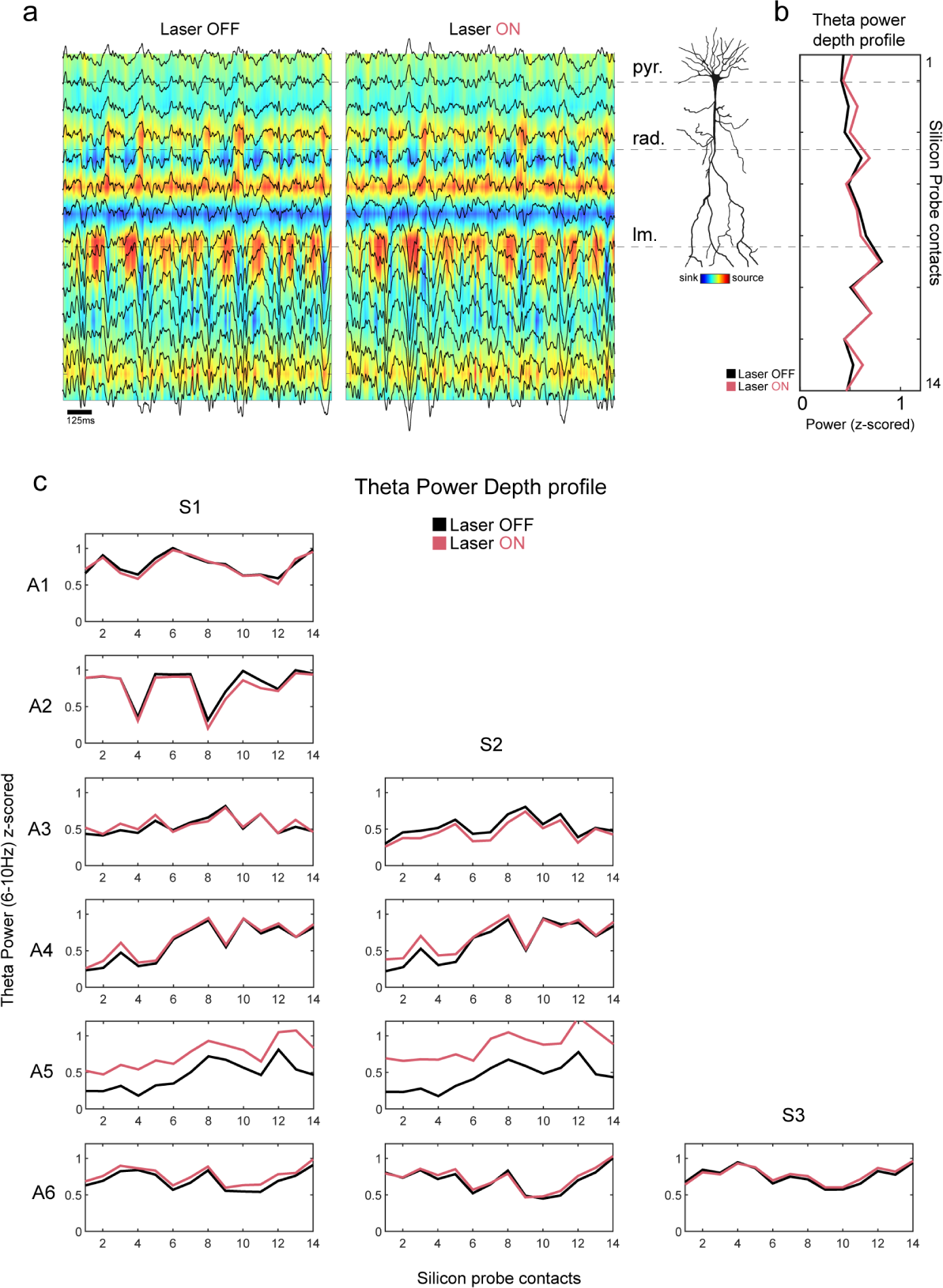
Layer-resolved theta power profiles (6-10 Hz) in CA1 during optogenetic inhibition of EC LIII direct input. (a) Representative examples of color coded sink and sources currents during laser OFF and laser ON periods. Local field potential traces are superimposed in black. Silicon probe contacts span all CA1 layers. (b) Average theta power depth profile, same session as in (a). (c) Theta power profiles across CA1 layers. Periods of immobility (movements *<* 3 cm/s) were removed. All animals (N=6), all session (n=11). No differences are found between conditions (laser OFF vs laser ON) - especially in the s.l.m., where EC LIII connections reach CA1 pyramidal neurons. Higher power profiles in both sessions of the Animal 5 are due to higher velocity performances during laser OFF and laser ON runs (Figure S2) - faster velocities coincide with higher power in hippocampal oscillations at all frequency bands [1].

**Fig. S12.**
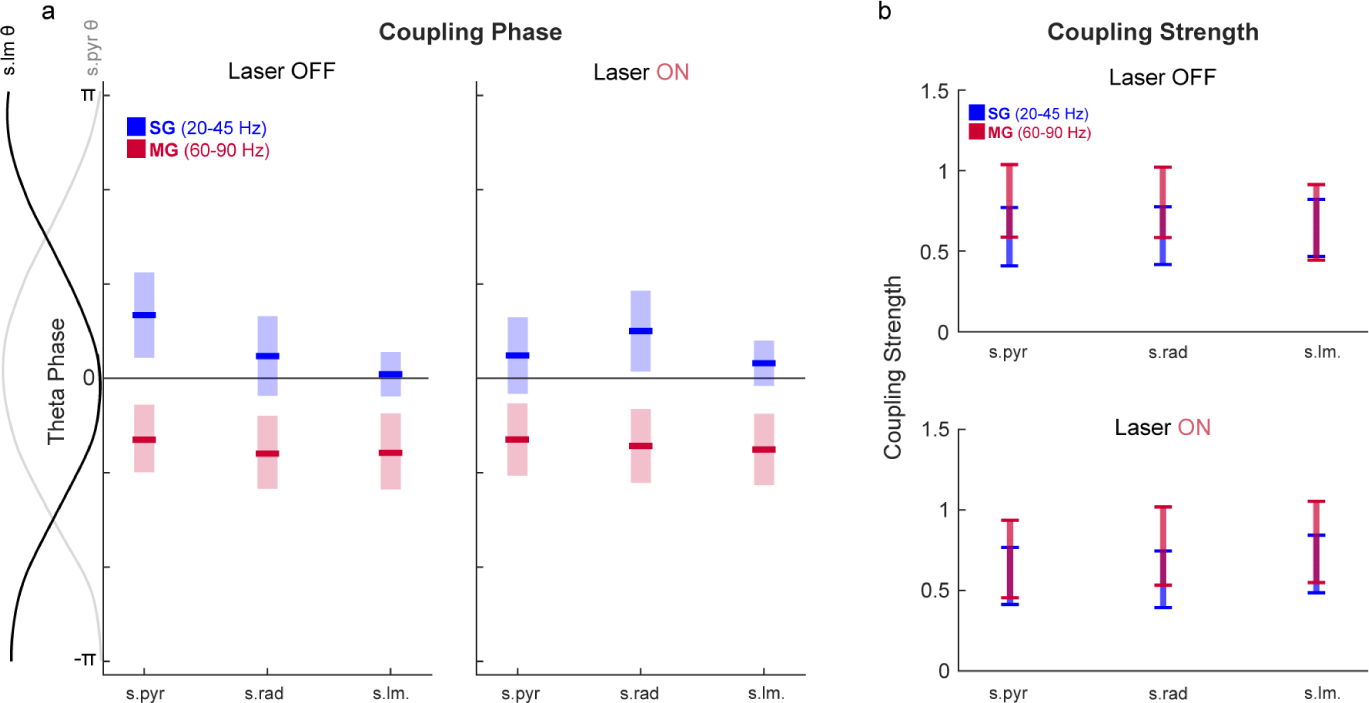
Theta-gamma coupling properties in CA1 during transient optogenetic inhibition of EC LIII direct input. (a) Coupling strength profiles of the slow and medium gamma components to the underling theta rhythm, across all CA1 layers. (b) Theta phase preference profiles of the slow and medium gamma components across all CA1 layers. No differences in both coupling strength and preferred coupling phase profiles. SG = slow gamma (20-45 Hz). MG = medium gamma (60-90 Hz).

